# Dynamics, cariogenicity, and geographic distribution of *Streptococcus mutans* adhesin types suggest adaptation to individual hosts

**DOI:** 10.1101/514265

**Authors:** Nongfei Sheng, Anders Esberg, Lena Mårell, Elisabeth Johansson, Carina Källestål, Nicklas Strömberg

**Affiliations:** Department of Odontology/Cariology and Oral health, Umeå University, SE-901 87 Umeå, Sweden; Department of Women’s and Children’s Health, Uppsala University, Uppsala, Sweden; Department of Disease Control, London School of Hygiene & Tropical Medicine, London, UK.

## Abstract

Dental caries, the most common chronic infectious disease, involve *Streptococcus mutans* SpaP A/B/C and Cnm/Cbm adhesin types of different virulence. We have explored their stability and dynamics over 5 years, their geographic distribution, as well as the potentially increased cariogenicity of specific SpaP B subtypes. We performed qPCR and TaqMan typing using whole saliva and isolates from 452 Swedish adolescents followed from 12 to 17 years of age. Approximately 50% of the children were infected at baseline with a single dominant (44%) or mixed (6%) SpaP A, B, or C type, some of which were also Cnm (6%) or Cbm (1%) positive. Stability (+, +) was high for *S. mutans* infection (85%) and dominant SpaP A or C (80% and 67%) and Cnm or Cbm (85% and 100%) types, but low for SpaP B (51%) and mixed SpaP A/B/C types (26%). Only five children switched from one SpaP type to another, and none between Cnm and Cbm types. Mixed SpaP A/B/C types were typically lost or changed into dominant types. Moreover, children infected with Cnm (n=26) types were more frequent in the northern (Skellefteå) region (p = 0.0041), and those with Cbm types (n=7) in the southern (Umeå) region. Children infected with SpaP B-2 subtypes had a doubled caries experience (p = 0.009) and 5-year caries increment (p = 0.02) compared to those infected with SpaP A. In conclusion, the stable dominant but instable mixed adhesin types and geographic differentiation of Cnm and Cbm types suggest adaptation of low and high cariogenicity types to specific individuals.

## Introduction

Dental caries affect billions of people, with large differences in prevalence and activity between populations and individuals [1–3]. In populations with a high prevalence of caries, the predominant cause of the disease is poor lifestyle in terms of frequent sugar intake and lack of oral hygiene, which promote acid-producing and acid-tolerant bacteria, such as *Streptococcus mutans* and non-mutans streptococci [1,4,5]. Thus, infection by *S. mutans* is a clinical risk factor that, together with the oral microbiome composition of acid-tolerant and acid-producing members, coincides with prospective caries development [5–7].

In Sweden and other populations that have shifted from a high to low prevalence of caries, the majority of children are either free or nearly free of caries and ~20% so-called high-risk children carry most of the disease burden [8,9]. The high-risk children develop more severe caries that are not prevented effectively by standard lifestyle and fluoride interventions [8,9]. Thus, today’s low prevalence population may involve subtypes of caries caused by genetic and previously unresolved factors [10]. Recently, *PRH1* and *PRH2* immunodeficiency in controlling moderate cariogenic oral non-mutans streptococci, or carriage of high cariogenicity types of *S. mutans* strains emerged as the primary causes of severe caries in 452 Swedish adolescents followed from 12 to 17 years of age [7,11,12]. Accordingly, children infected with *S. mutans* SpaP B and Cnm adhesin types developed more caries than those infected with SpaP A or C and Cbm types or non-infected children [7]. The Cnm phenotype, which occurs in 6% of adolescents, has been implicated in endocarditis and stroke [13–15]. Both Cnm and SpaP B, one of the three core genome SpaP A, B, or C adhesin types in *S. mutans* biotypes A, B, and C, exhibit increased acid tolerance and binding to saliva pattern recognition receptor DMBT1 [7,16,17]. Moreover, sequence typing of 70 caries-free and decayed extreme cases suggested a cluster or subgroup of SpaP B-2 types with increased cariogenicity [7]. However, the high cariogenic nature of SpaP B-2 types remains to be verified in the entire sample of 452 adolescents.

*S. mutans* infects 40-80% of subjects and is transmitted similarly to the oral and gut microbiomes from parent to child [18,19]. Infection by *S. mutans* is dominated by one or a few phenotypes, such as the SpaP A, B, or C adhesin types [18,19], whereas genotype diversity in oral streptococci increases with extent of sequencing [20]. The *S. mutans* adhesin types and oral, gut, and other microbiomes colonize their niches in a stable fashion over time, though quantitative fluctuations occur in response to antibiotic treatment, traveling, breast feeding, and unknown factors [7,21–23]. However, whether *S. mutans* adapts to host receptor repertoires similar to uropathogenic *Escherichia coli* [24], *Helicobacter pylori* [25], and Noro/HIV viruses [26,27] upon infection in individuals, ethnic groups, or animal species is unknown [28]. The stability and dynamics over time and geographic differentiation of *S. mutans* adhesin types may shed light on their adaptation to specific individual hosts.

The aims of this study were to explore the stability, dynamics, and geographic differentiation of infection by *S. mutans* adhesin types during adolescence and puberty, and to verify the high cariogenic nature of SpaP B-2 subtypes in the entire sample of 452 Swedish adolescents followed from 12 to 17 years of age.

## Material and methods

### Study sample of 452 adolescents

A total of 452 children were enrolled from 13 Public Dental Service Clinics in northern Sweden as caries cases-controls based on caries data and examined at 12 and 17 years of age for caries (Decayed, enamel included, Filled Surfaces, DeFS), lifestyle, and biological variables [7,11]. The children received operative treatment and caries prevention according to the ordinary routines and policies of the clinics. The exclusion criterion was unwillingness to participate in the study. Of the 452 children, 390 were re-examined at 17 years of age, resulting in a 14% dropout rate (62/452) for having moved out of the area (n=20) or repeatedly missing the examination (n=42). The ethics committee for human experiments at Umeå University, Sweden, approved the study and all parents signed informed consent to participate.

### Whole saliva sampling and preparation of DNA

Whole saliva was collected from the children at 12 and 17 years of age by chewing on paraffin. One child failed to produce saliva at 12 years of age, and another at 17 years of age. DNA was prepared from 400 μl of each saliva sample using the GenElute ™ Bacterial Genomic DNA Kit (Sigma Aldrich, Sweden) [7].

### Typing of *S. mutans* infection by quantitative polymerase chain reaction (qPCR)

Infection by *S. mutans* was typed by qPCR using pure DNA from whole saliva, the KAPA SYBR FAST Universal qPCR kit (Sigma-Aldrich, Sweden), and primers specific to the conserved *gtfB* and *gtfC* regions in a Corbett Rotor-Gene 6000 apparatus [29] (S1 Table). Quantitative calibration curves used DNA purified from serial dilutions of an *S. mutans* reference strain [7], and the cut-off value for *S. mutans* positive versus negative saliva was 3000 pg.

### Genotyping of *spaP* and *cnm/cbm* adhesin types by qPCR

Adhesin types in whole saliva were genotyped by qPCR using pure DNA from whole saliva, the KAPA SYBR Fast Universal qPCR kit (Tectum, Sweden), and primers specific to each adhesin type in a Corbett Rotor gene 600 apparatus [7] (S1 Table). The primers for *cnm* and *cbm* did not cross-react between or with other templates, and the primers for *spaP A*, *B*, and *C* were from the *spaP* sequences based on prior testing and lack of cross-reactivity between *A*, *B*, and *C* or with DNA from oral streptococci with *spaP* analogs. The typing used internal standards and quantitative calibration curves from dilutions of DNA purified from a reference genotype of each adhesin type and cut-off values for *A* (3000 pg), *B* (3000 pg), *C* (6000 pg), *cnm* (3000 pg), and *cbm* (1000 pg).

### *S mutans* strains from infected children

The *S. mutans* strains were isolated from plaques from infected children, characterized, and typed for SpaP A/B/C and Cnm/Cbm status as described previously [7]. One strain representative of the dominant SpaP A, B, or C type in the saliva of each of 214 of the 217 infected children was used. DNA was prepared from a 3 ml culture of the representative strain using the GenElute ™ Bacterial Genomic DNA Kit (Sigma Aldrich, Sweden) [7].

### Genotyping of children as positive or negative for SpaP B-2 subtypes

Children were genotyped positive or negative for infection with *S. mutans* SpaP B-2 subtypes by genotyping their whole saliva as positive for *spaP A*, *B,* or *C,* followed by TaqMan typing of DNA from one representative *S. mutans* isolate for B-2 status using canonical SNPs [7] (S1 Table). The following canonical SNPs specific to *spaP* subtypes B-2 and *spaP A*, *B,* and *C* were selected and designed using MEGA6 software [30] and evaluated by the Thermofisher custom genomic TaqMan assay and design pipeline for non-human templates (https://www.thermofisher.com/order/custom-genomic-products/tools/genotyping) and analyzed using the ABIprism 7900HT (Applied Biosystems): SpaP SNP_2137-2139_ [GT/CG], where [GT] defines SpaP A and [CG] defines SpaP B and C type; SpaP_3188_ [A/G], where [A] defines SpaP AB-type and [G] defines SpaP C; SpaP_3045_ [A/G], where [A] defines B_2_ and [G] defines non-B_2_. All TaqMan assays were performed using the TaqMan™ Genotyping Master Mix (Applied Biosystems™) and the predesigned PCR program consisting of an initial denaturing at 95°C for 10 min, followed by 40 cycles of 92°C for 15 sec. and 60°C for 1 min. TaqMan typing of the 70 reference isolates for SpaP A/B/C and SpaP B-2 subtypes validated the canonical SNPs and recalled the expected sequence types except for one isolate.

### Geomapping of *S. mutans* genotypes

Using Google maps, coordinates for each individual residential address with SpaP A, B, or C including B-2 and Cnm/Cbm isolates were plotted on a map. The division allocated the North and South regions. The two regions were divided into city and countryside areas.

### Data analysis

Data were expressed as means ± standard deviations (SDs). The statistical analyses were performed using Mann-Whitney U tests and chi^2^ or Fisher exact tests in Rcmdr or SPSS software. All statistical analyses used two-tailed tests, and p-values < 0.05 were considered significant.

## Results

### Dynamics of *S. mutans* adhesin types during adolescence

To explore the dynamics of infection with *S. mutans* and its SpaP A/B/C and Cnm/Cbm adhesin types during adolescence, we typed *S. mutans* in whole saliva collected from 452 Swedish adolescents at 12 and 17 years of age using qPCR (Table 1, S1 Figure). Forty-eight percent of the children were infected with *S. mutans* at 12 years of age according to *gtf* typing, and infection increased to 55% at 17 years of age (p =0.033). The stability of children infected (+ to +) or not infected (− to −) over the 5-year period was 85% and 71%, respectively, with loss (+ to −) and more frequently gain (− to +) of infection occurring in every forth to sixth child (Table 2).

**Table 1.** Prevalence of *S. mutans* SpaP and Cnm/Cbm adhesin types in Swedish adolescents.

| Type<sup>a</sup> | Prevalence<sup>b</sup> |  |  |  | *p*<sup>c</sup> |
| --- | --- | --- | --- | --- | --- |
|  | 12 years old (N=452) |  | 17 years old (N=390) |  |  |
|  | n | % | n | % |  |
| *S. mutans* | 217 | 48 | 216 | 55 | 0.033 |
| Core genome |  |  |  |  |  |
| SpaP A | 111 | 24 | 113 | 29 | 0.148 |
| SpaP B | 71 | 16 | 39 | 10 | 0.014 |
| SpaP C | 19 | 4 | 24 | 6 | 0.212 |
| Mixed SpaP A, B, or C | 27 | 6 | 33 | 9 | 0.162 |
| Pan genome |  |  |  |  |  |
| Cnm | 26 | 6 | 19 | 5 | 0.646 |
| Cbm | 7 | 1.5 | 6 | 2 | 1 |

<sup>a</sup> *S. mutans* types measured in whole saliva by qPCR in adolescents at 12 and 17 years of age.
<sup>b</sup> Prevalence in adolescents at 12 and 17 years of age. Data were collected from 452 children at 12 years old and from 390 adolescents at 17 years old.
<sup>c</sup> p-value from $\chi^2$ test.

**Table 2.**
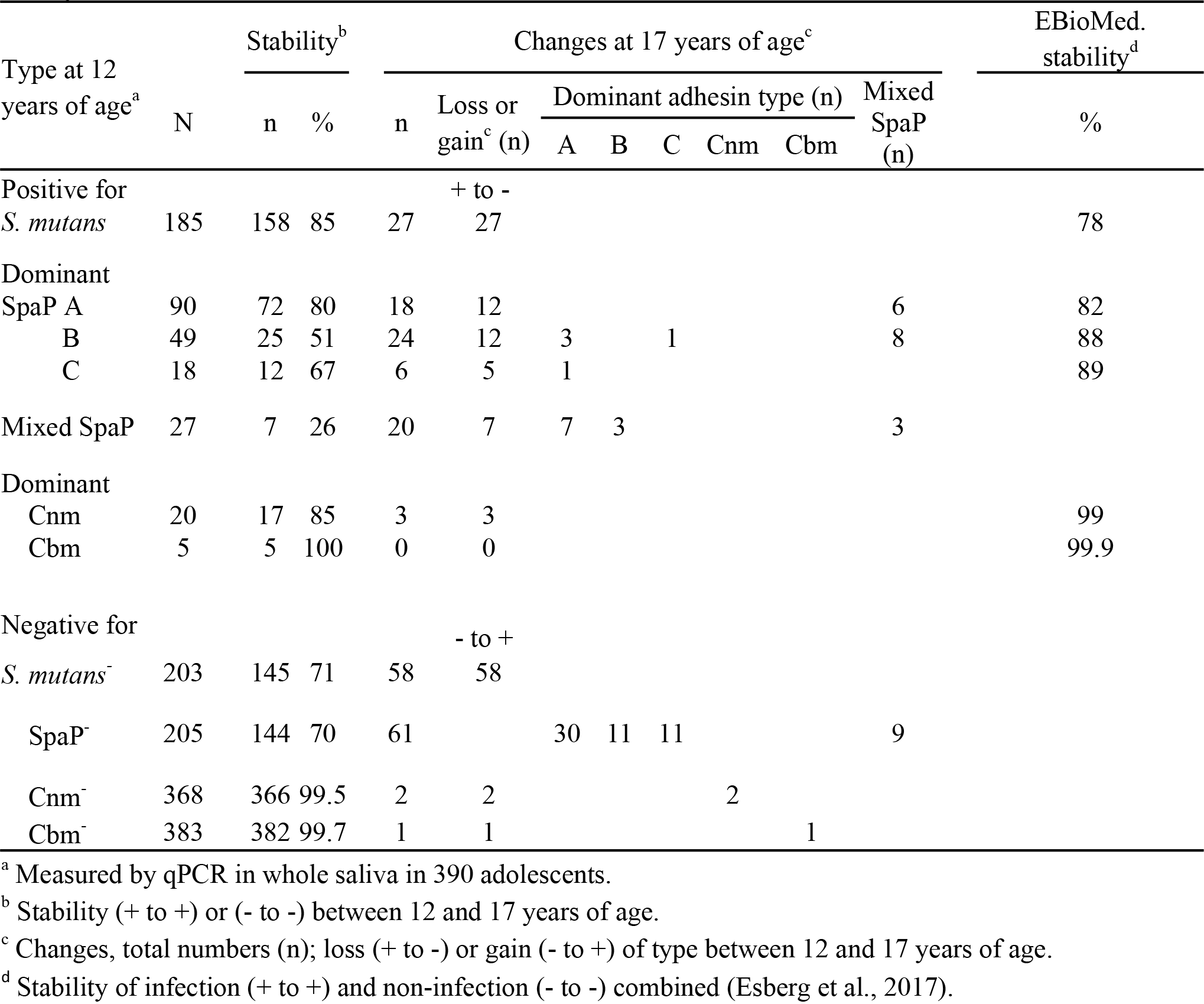
Stability and dynamics of SpaP and Cnm/Cbm adhesin types during adolescence in Swedish 12 to 17 year olds.

Half of the infected children carried either a single dominant SpaP A, B, or C (44%) or mixed SpaP A, B, or C (6 %) adhesin type according to *spaP* typing. Some children were also positive for Cnm (6%) or Cbm (1.5%, Table 1). The stability (+ to +) of single dominant SpaP A/B/C and Cnm/Cbm types was 80/51/67% and 85/100%, respectively, with only five children switching from one dominant SpaP A, B, or C types to another, and none between Cnm or Cbm type (Table 2, S2 Figure). The prevalence of infection with SpaP B of low stability (+ to +, 51%) decreased from 16% to 10 % over the 5-year period (p = 0.014, Table 1), and SpaP B was more frequently lost or transformed into dominant SpaP A C (n=4) or mixed types than SpaP A, which only transformed into mixed types (Table 2).

In contrast, the stability (+ to +) of mixed SpaP A/B/C types was low (26%, Table 2). Accordingly, they were either lost or transformed into single dominant types over time, and non-infected children less frequently transformed into mixed types (14% or 9/61) than dominant types (96% or 52/61, Table 2).

Quantitative numbers of *S. mutans* and its types fluctuated between infected children, as well as between 12 and 17 years of age, as shown for the SpaP B type (Fig. 1). However, distinct changes were less frequent in non-infected children than infected children with exponentially higher numbers of the organism.

**Fig 1.**
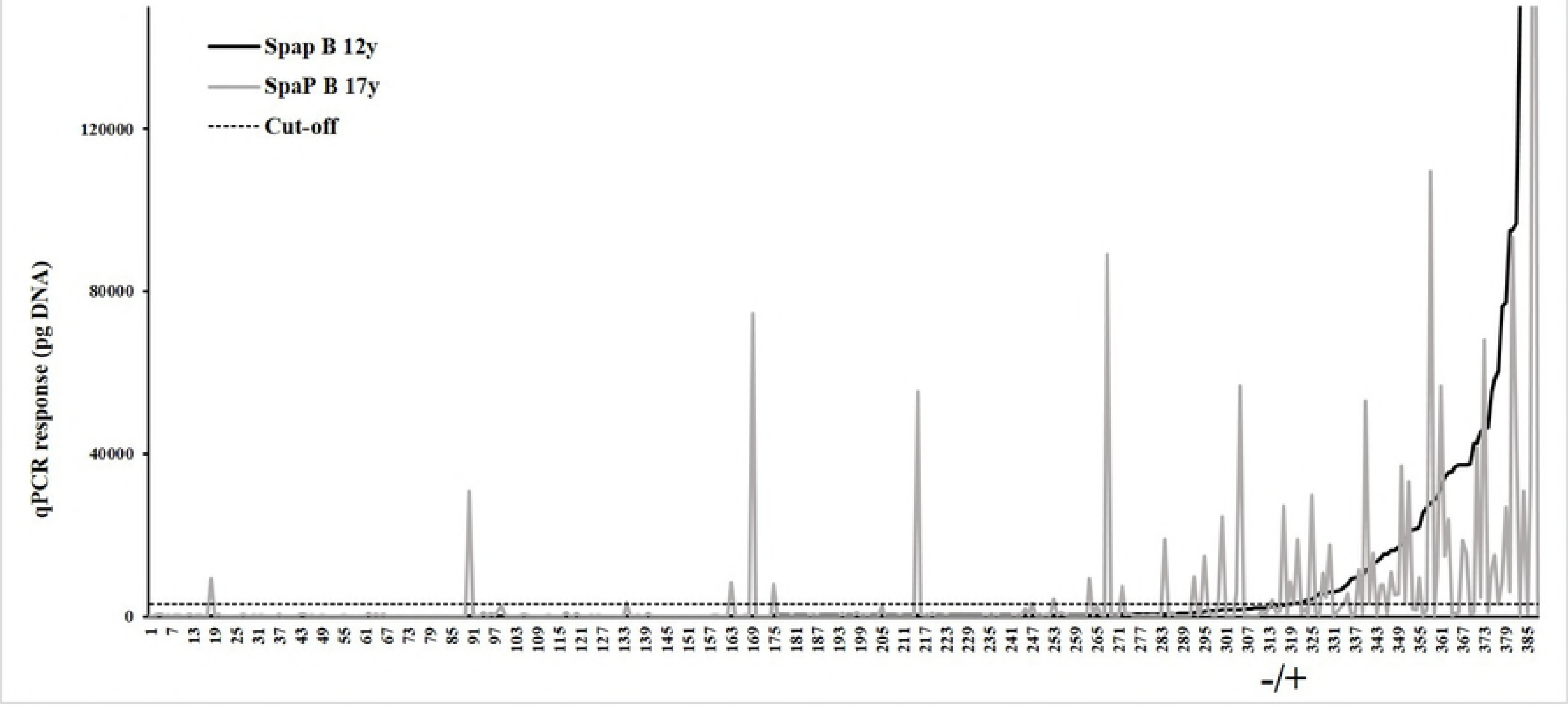
Range of *S. mutans* SpaP B in infected children at 12 years of age (bold line) and changes at 17 years of age (grey line). Quantitative numbers of *S. mutans* SpaP B are given as the qPCR response (pg DNA) in each child at 12 (bold line) and 17 (grey line) years of age. The cut-off level (dotted line) marks infected and non-infected children at 12 years of age (− / +).

### Geographic differentiation of children infected with Cnm and Cbm types

Next, we explored how children infected with SpaP A/B/C and Cnm/Cbm types were distributed over the southern (Umeå) and northern (Skellefteå) regions of the county of Västerbotten in Northern Sweden (Fig. 2, S2 Table). Children infected with SpaP A/B/C or infected versus non-infected children were distributed evenly between the two regions, whereas those with Cnm types (n=26) appeared more frequently in the northern (Skellefteå) region (p = 0.0041) and Cbm types (n=7) in the southern (Umeå) region.

**Fig 2.**
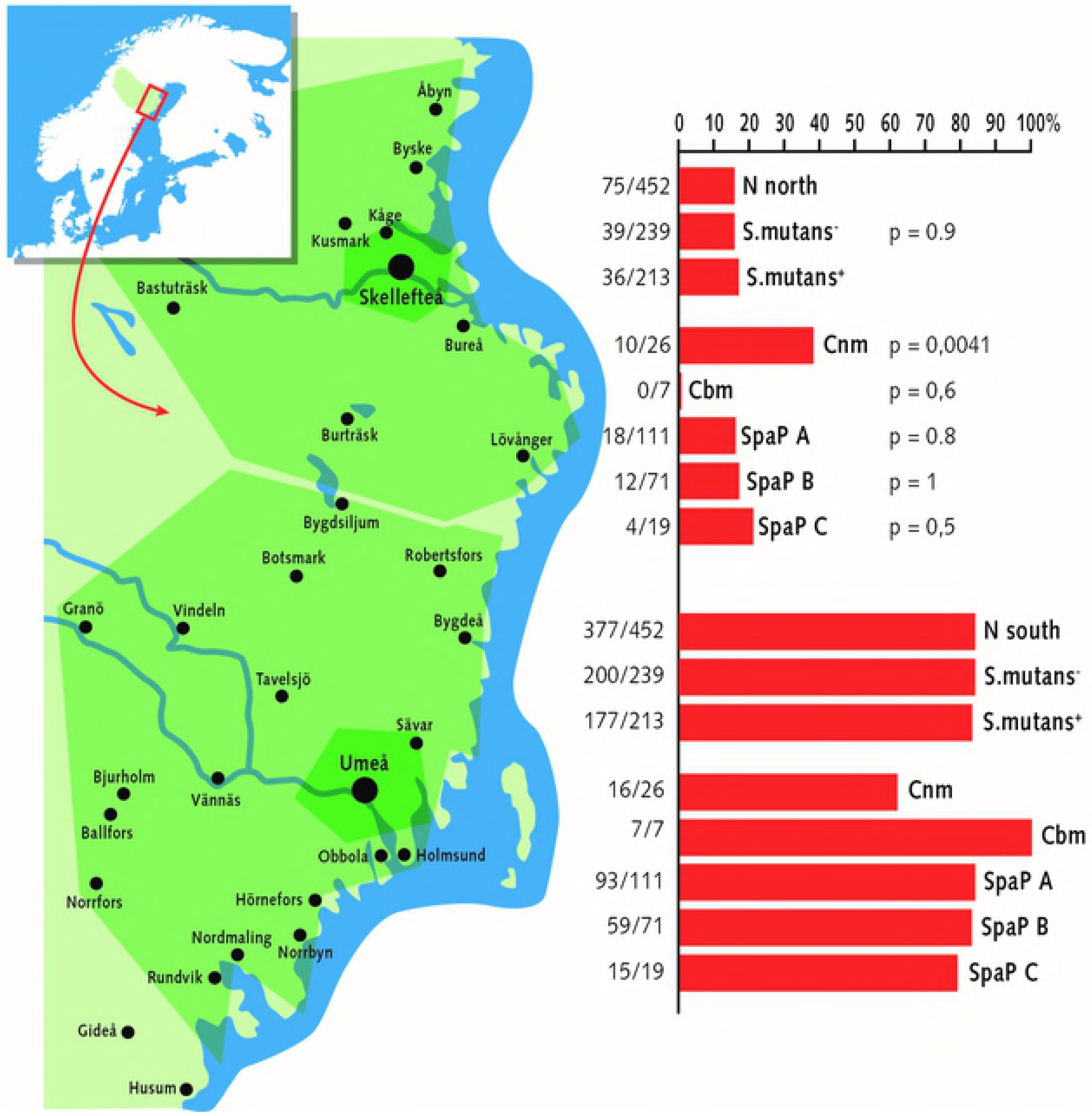
Different relative distributions of the Cnm and Cbm adhesin types in the south (Umeå) and north (Skellefteå) regions of northern Sweden. Bars indicate the relative proportion (%) of children with dominant SpaP A, B, or C and Cnm or Cnm adhesin types in each south and north region at 12 years of age. N = total numbers of children sampled in each region. The numbers to the left of each bar is the number of adhesin types in the region out of the total number in the county (more children were sampled in the south). The p-value to the right is the significance of the difference in distribution of each type between the two regions.

### Verification of high cariogenicity SpaP B-2 subtypes

Sequencing of isolates of *S. mutans* from 70 decayed and caries-free extreme cases in the 452 adolescents suggested a cluster of SpaP B-2 subtypes of increased cariogenicity [7]. To verify the high cariogenic nature of the SpaP B-2 types, children with saliva positive for SpaP A, B, or C based on qPCR typing were typed for the B-2 status of representative isolates of *S. mutans* by TaqMan (Table 3). Children infected with SpaP B-2 subtypes had a doubled baseline caries (p = 0.009) and 5-year caries increment (p = 0.02) compared to children infected with the SpaP A type (Table 3).

**Table 3.** Verification of high cariogenicity of *S. mutans* SpaP B-2 in Swedish adolescents.

| Type<sup>a</sup> | Type | DeFS-12y<sup>b</sup> | | | DeFS-17y<sup>b</sup> | | | ΔDeFS-5y<sup>c</sup> | | |
| --- | --- | --- | --- | --- | --- | --- | --- | --- | --- | --- |
| | | mean±SD | n | $p^d$ | mean±SD | n | $p^d$ | mean±SD | n | $p^d$ |
| Positive | | 3.6 ± 3.0 | 217 | $8 \times 10^{-13}$ | 9.2 ± 8.8 | 185 | $4 \times 10^{-8}$ | 5.5 ± 7.1 | 185 | $3 \times 10^{-5}$ |
| Negative | | 1.7 ± 2.1 | 234 | Ref. | 4.9 ± 5.7 | 204 | Ref. | 3.1 ± 4.9 | 204 | Ref. |
| SpaP | A | 3.0 ± 2.8 | 111 | Ref. | 8.4 ± 8.2 | 96 | Ref. | 5.2 ± 7.1 | 96 | Ref. |
| | B | 4.2 ± 3.5 | 71 | 0.034 | 11.4 ± 11.1 | 55 | 0.045 | 6.7 ± 8.8 | 55 | 0.110 |
| | C | 3.3 ± 1.8 | 19 | 0.365 | 7.5 ± 4.8 | 18 | 0.746 | 4.3 ± 4.1 | 18 | 0.929 |
| SpaP B<sup>c</sup> | B-2 | 5.5 ± 3.4 | 13 | 0.009 | 16.5 ± 11.8 | 8 | 0.013 | 11.9 ± 11.9 | 8 | 0.020 |
| | A | 3.0 ± 2.8 | 111 | Ref. | 8.4 ± 8.2 | 96 | Ref. | 5.2 ± 7.1 | 96 | Ref. |

<sup>a</sup> Children infected with *S. mutans* (positive) or not (negative) and SpaP subtypes as measured by qPCR and TaqMan in 452 children at 12 years of age.
<sup>b</sup> Caries DeFS (decayed, enamel included, filled surfaces) at 12 and 17 years of age.
<sup>c</sup> $\Delta$ DeFS (5y) = 5-year prospective caries increase from 12 to 17 years of age.
<sup>d</sup> Two-sided p-value from Mann-Whitney U-test.
<sup>e</sup> B-2 marks children infected with SpaP B-2 subtype.

## Discussion

The dynamics and geographic differentiation of infection with *S. mutans* adhesin types in 452 Swedish adolescents from 12 to 17 years of age suggest their adaptation to specific individual hosts. We recently used the same study sample to identify subtypes of caries caused primarily by lifestyle, immunodeficiency, or high cariogenicity type *S. mutans* [7,11]. This knowledge will facilitate new models for investigating the etiology, prevention, and treatment of caries and further resolve the role of *S. mutans* and oral streptococci in caries development. Severe caries cases may fail to control the commensal non-mutans streptococcal flora (*i.e.*, immunodeficiency type caries) or carry high cariogenicity type *S. mutans*, whereas infection by *S. mutans* seems to promote the lifestyle type caries [7,11].

We found an overall high infection stability for *S. mutans* (85%), and its prevalence increased from 48% to 55% over 5 years (p = 0.033). Consequently, roughly four out of five children are stably infected or non-infected, and about every fifth child shows a loss or gain of infection by *S. mutans* and by type. The prevalence of SpaP B with increased cariogenicity decreased from 16% to 10% over 5 years (p = 0.014). This decline may arise from the low stability (51%) and gain of infection (14%) by SpaP B and the treatment, including extra fluorides, given to high caries cases. Together with the consumption of caries-sensitive sites with time, this may contribute to explaining the attenuation of caries activity in later adolescence [31]. The decline in SpaP B may also involve molecular changes induced by puberty, such as glycosylation of DMBT1 receptors for *S. mutans*, as suggested by the inverse association of saliva-binding *S. mutans* with caries at 12 and 17 years of age [7]. However, the quantitative numbers of *S. mutans* types fluctuated markedly, despite the stability in terms of quality (+ or −) and associated caries development [7], and should be interpreted with caution in terms of plaque quality and virulence.

The stable nature of dominant versus mixed adhesin types of *S. mutans* suggests strong selective and adaptive forces at the individual level. The stability (+ to +) of 51% to 100% for dominant SpaP A, B, or C and Cnm or Cbm types is consistent with their reported stability of 78% to 100% for infection (+ to +) and non-infection (− to −) status combined [7]. The low stability (26%) of mixed SpaP A/B/C and absence of Cnm/Cbm types, as well as the cross-sectional predominance of dominant over mixed types and in gain (− to +) of infection, suggest that mixed types easily transform into dominant types. The fact that only five children switched from one SpaP to another, and none between Cnm and Cbm type, further emphasizes the inherent stability of dominant types. Similar to recent microbiome studies, our findings suggest that bacterial colonization is stable in terms of quality and that switching *S. mutans* adhesin type is a rare and unstable event. However, similar to carriage of *Streptococcus mitis* [20], the adolescents may carry a mixture of dominant and minor *S. mutans* adhesin types that fluctuate in number over time. The environmental and host genetic factors or receptor mechanism for adaptation of dominant SpaP A/B/C and Cnm/Cbm types to specific individuals is not yet known.

The present findings show a geographic differentiation in infection, with Cnm (n = 26) and Cbm (n = 7) types being more frequent in the north (Skellefteå, p = 0.0041) and south (Umeå, 7 out of 7 children) region, respectively. However, since the more prevalent SpaP A and B types as well as infected versus non-infected children distributed evenly between the two regions, both prevalence and biological factors may contribute to the observed differences. The transmission of *S. mutans* from parents to child and low prevalence will inherently generate localized family patterns in regions of low mobility. Unknown environmental, host genetic, and ethnic factors may also influence the Cnm/Cbm and SpaP types of core and pan genome origin, respectively. Another oral commensal pathogen, *Aggregatibacter actinomycetemcomitans* clone JP2, associated with juvenile periodontitis, transmits among African Americans for cultural or host genetic reasons [32], and residents along the river valleys in northern Sweden exhibit distinct genetic repertoires [33], with familial amyloid polyneuropathies markedly increased in the Skellefteå region [34]. Therefore, infection with SpaP A/B/C and Cnm/Cbm adhesin types should be further investigated in terms of family patterning and host receptor repertoires [11,12].

We verified the high cariogenicity of SpaP B-2 subtypes by genotyping saliva and representative *S. mutans* isolates from all 452 adolescents. Children with SpaP B-positive saliva and B-2 isolates had significantly increased caries experience and prospective 5-year caries increase compared to children infected with SpaP A types. The cluster of B-2 isolates with unique mutations in *spaP* and housekeeping genes for saliva binding and acid tolerance, respectively, may account for a large portion of the cariogenicity of *S. mutans* biotype B [7]. Although the present SpaP B-2 typing scheme extends our previous sequencing of isolates from a few to all 452 adolescents, it uses only a representative isolate of *S. mutans* from each child. Further evaluation of the relative cariogenicity of *S. mutans* adhesin types should be performed in prospective studies with sequence typing of saliva, as well as multiple isolates in both the primary and permanent dentitions.

The overall stable but in every fifth adolescent changing infection with *S. mutans* and its adhesin types is consistent with the persistent nature of caries and important for microbial diagnostic and therapeutic approaches. This finding implies biomarkers for both risk and treatment, such as use of chlorhexidine or other anti-microbial treatments. However, greater understanding of the colonization and persistence factors will be necessary for therapies, such as future replacement of high virulence types with low virulence or probiotic types. We hypothesize that children without caries and those not infected with *S. mutans* may carry a healthy microbiome with universal core members that can be used in future probiotic cocktails to treat or prevent caries [21]. *S. mutans* is transmitted from parent to child during the perinatal and infancy period, when the microbiome, nutrition, and immune system synergize to shape oral tolerance, the innate and adaptive immunity, and the human phenotype [35,36]. Such early encounters between SpaP A/B/C and Cnm/Cbm types and the host innate and adaptive immunity may shape distinct long-lasting immunity and inflammation responses. Thus, the perinatal and infancy period may represent a window of opportunity for establishing long-lasting conditions beneficial to oral and general health [37], and the association of Cnm types and tooth loss with stroke and systemic risk further emphasizes the potential of exploring *S. mutans* phenotypes in the prevention of caries early in life.

## Acknowledgment

Financial support for this study was received from a regional agreement between Umeå University and Västerbotten County Council in the field of Medicine, Odontology, and Health (377341); a Fund for Cutting-Edge Medical Research grant from the County Council of Västerbotten (135041); and from the Swedish Dental Society. We also acknowledge local foundations at Umeå University. We thank Ingmarie Bernhardsson, Ulla Öhman, Ewa Strömqvist-Engbo, and Rolf Claesson for assistance, and the families and public health care personnel who participated in the study.

## Supporting Information

**S1 Figure. Representative examples of typing SpaP A/B/C adhesin types and major changes in the infection pattern between 12 and 17 years of age.** Bars show the qPCR response (pg DNA) when genotyping whole saliva from adolescents at 12 years and 17 years of age (12y and 17y, respectively). The cut-off level distinguishes positive and negative adhesin types. The mean ± SD response for dominant SpaP A, B, and C types at baseline are 52864±79777, 39907±56161, and 39193±57668, respectively. The major changes are gain (− to +) and loss (+ to −) of adhesin type, mixed to single type or vice versa, and switch from one type to another. (TIF)

**S2 Figure. The five children that switched from one single dominant SpaP A, B, or C adhesin type to another type between 12 and 17 years of age.** Bars show the qPCR response (pg DNA) when genotyping whole saliva from adolescents at 12 and 17 years of age (12y and 17y, respectively). The cut-off level (dotted line) marks infected versus non-infected status. (TIF)

**S1 Table. Primers and SNPs for SpaP and Cnm/Cbm genotyping by qPCR and Taqman.** (PPTX)

**S2 Table. Geographic distribution of** *S. mutans* **adhesin types in Swedish adolescence at 12 years of age in Västerbotten County, Sweden.** (PPTX)

## Author Contributions

Conceptualization: Nongfei S, Nicklas S.
Data curation: Nongfei S, Anders E, Nicklas S.
Formal analysis: Nongfei S, Nicklas S.
Funding acquisition: Nongfei S, Carina K, Nicklas S.
Investigation: Nongfei S, Anders E, Lena M, Elisabeth J, Carina K, Nicklas S.
Methodology: Nongfei S, Anders E, Lena M, Elisabeth J, Carina K, Nicklas S.
Project administration: Carina K, Nicklas S.
Resources: Carina K, Nicklas S.
Software: Nongfei S, Anders E, Carina K, Nicklas S.
Supervision: Carina K, Nicklas S.
Validation: Anders E, Carina K, Nicklas S.
Visualization: Nongfei S, Anders E, Lena M, Elisabeth J, Carina K, Nicklas S.
Writing – original draft: Nongfei S, Nicklas S.

